# Cysteine: an ancestral Cu binding ligand in green algae?

**DOI:** 10.1101/2023.03.15.532757

**Authors:** Daniela Strenkert, Stefan Schmollinger, Yuntao Hu, Christian Hofmann, Kristen Holbrook, Helen W. Liu, Samuel O. Purvine, Carrie D. Nicora, Si Chen, Mary S. Lipton, Trent R. Northen, Stephan Clemens, Sabeeha S. Merchant

## Abstract

Growth of *Chlamydomonas reinhardtii* in zinc (Zn) limited medium leads to disruption of copper (Cu) homeostasis, resulting in up to 40-fold Cu over-accumulation relative to its typical Cu quota. We show that Chlamydomonas controls its Cu quota by balancing Cu import and export, which is disrupted in a Zn deficient cell, thus establishing a mechanistic connection between Cu and Zn homeostasis. Transcriptomics, proteomics and elemental profiling revealed that Zn-limited Chlamydomonas cells up-regulate a subset of genes encoding “first responder” proteins involved in sulfur (S) assimilation and consequently accumulate more intracellular S, which is incorporated into L-cysteine, γ-glutamylcysteine and homocysteine. Most prominently, in the absence of Zn, free L-cysteine is increased ~80-fold, corresponding to ~ 2.8 × 10^9^ molecules/cell. Interestingly, classic S-containing metal binding ligands like glutathione and phytochelatins do not increase. X-ray fluorescence microscopy showed foci of S accumulation in Zn-limited cells that co-localize with Cu, phosphorus and calcium, consistent with Cu-thiol complexes in the acidocalcisome, the site of Cu(I) accumulation. Notably, cells that have been previously starved for Cu do not accumulate S or Cys, causally connecting cysteine synthesis with Cu accumulation. We suggest that cysteine is an *in vivo* Cu(I) ligand, perhaps ancestral, that buffers cytosolic Cu.

## INTRODUCTION

First row transition metals are essential for life. As cofactors in enzymes, they enable reactions that cannot readily be catalyzed by organic molecules. Among the transition elements, Cu is a key player in redox chemistry and reactions involving oxygen chemistry or oxygenated substrates (Merchant et al., 2020). Cu entered biology well after Fe, as its solubility in the aquatic milieu that supports life increased only after oxygenation of the planet, giving rise to new bioenergetic capabilities like aerobic respiration (Crichton and Pierre, 2001). Unsurprisingly, Cu enzymes are found in all aerobic organisms, making Cu essential in most habitats. Yet, its reactivity, especially in the presence of oxygen, renders it toxic (Goldstein and Czapski, 1986). Cu excess is also a problem in biological systems because of its ability to form stable Cu(I) or Cu(II) complexes with functional groups on biological macromolecules (Tottey et al., 2012). Therefore, organisms in all kingdoms of life have evolved mechanisms to handle Cu and control its delivery. The metabolism of Cu (defined as uptake, delivery, distribution and metalation of its binding site in proteins) is tightly regulated so that organisms take up only the Cu that they need as metal cofactors (Peña et al., 1999). Because of this, the Cu content of a cell closely matches the Cu-protein content. The abundance of individual cuproproteins is determined by the cellular demand for the pathways in which they function. For example, in yeast and Chlamydomonas, a multicopper oxidase, required for high-affinity Fe uptake, is induced under Fe-deficiency, resulting in increased demand for Cu in low-Fe environments and a draw on intracellular Cu. Accordingly, CTRs and ATX1, involved in Cu assimilation and distribution are up-regulated (Dancis et al., 1994; La Fontaine et al., 2002). In situations where there is increased demand for aerobic respiration, mitochondrial cytochrome oxidase (a Cu-containing enzyme) is induced, and again, Cu chaperones are expressed to accommodate the higher intracellular Cu demand and facilitate Cu delivery to the mitochondria (Kropat et al., 2015). Transporters of the CTR type and P-type ATPases are key to maintenance of the Cu quota, which they achieve by balancing import and export, respectively (Dancis et al., 1994; Stangeland et al., 1997; Peña et al., 2000; Axelsen and Palmgren, 2001;Sancenón et al., 2003).

Besides assimilation and efflux, intracellular sequestration in specialized, acidic vacuoles (lysosome related organelles or acidocalcisomes) is another important component of Cu homeostasis, especially in situations of excess (Raguzzi et al., 1988; Szczypka et al., 1997;Urbanowski and Piper, 1999; Rees et al., 2004; Docampo et al., 2005; Simm et al., 2007; Roh et al., 2012; Hong-Hermesdorf et al., 2014; Tsednee et al., 2019). Sequestration serves a detoxification role, but also a storage role, which can provide a competitive advantage in subsequent periods of deficiency because the store can be re-mobilized (Thomine et al., 2003;Rees et al., 2004; Rees and Thiele, 2007; Hong-Hermesdorf et al., 2014).

For metals of interest in biology, but especially for Cu species, the ions that are not associated with proteins are typically in complex with low molecular weight ligands. Metallothioneins (MTs) and phytochelatins (PCs) are examples of metal binding ligands that provide protection against metal toxicity (Foster and Robinson, 2011). MTs are genome-encoded polypeptides found in animals, fungi, plants and some algae, but not in Chlamydomonas (Robinson, 1989; Rauser,1999). PCs are a group of metal-binding peptides with the general structure (γ-Glu-Cys)_n_-Gly (n = 2–11) (Grill et al., 1985; Cobbett, 2000) that are non-ribosomally synthesized from glutathione (GSH) by the constitutively expressed enzyme called PC synthase (Grill et al., 1985, 1987; Grill et al., 1989; Zenk, 1996; Ha et al., 1999; Cobbett, 2000; Schat et al., 2002). PCs are found in land plants, algae (including Chlamydomonas) and fungi (Kondo et al., 1985; Gekeler et al., 1988;Mutoh and Hayashi, 1988; Howe and Merchant, 1992; Kneer et al., 1992; Bräutigam et al., 2010;Scheidegger et al., 2011): they can bind many different metals, including Cu, and hence are important in metal overload situations (Rauser, 1999; Clemens, 2001).

*Chlamydomonas reinhardtii* is a valuable reference organism for the study of metal homeostasis in photosynthetic organisms (Merchant et al., 2006). Under standard laboratory growth conditions, Chlamydomonas maintains a Cu quota between 1-2.5 × 10^7^ Cu atoms / cell that is mainly determined by the abundance of only two cupro-proteins, plastocyanin in photosynthesis and cytochrome (Cyt) oxidase in respiration (Kropat et al., 2015). If Cu becomes limiting, Chlamydomonas can reduce its Cu quota by substituting the abundant chloroplast-localized plastocyanin with a functionally equivalent Fe-containing protein, cytochrome *c*_6_ (Merchant and Bogorad, 1986). The tight control of the Cu quota is disturbed only in Zn-deficient cells, where massive Cu over-accumulation occurs (Malasarn et al., 2013; Hong-Hermesdorf et al., 2014). The mechanism underlying disturbed Cu homeostasis is not known. The simple explanation of unintentional, non-specific Cu(II) entry on an induced Zn(II) transporter (ZRTs) was ruled out because ectopic expression of ZRTs did not cause Cu over-accumulation (Sommer et al., 2010). The location of the excess, intracellular Cu was visualized by various complementary microscopy techniques and localized to acidocalcisomes (lysosome-related organelles) where they are inaccessible for cuproprotein biosynthesis or for turning off the nutritional Cu regulon, resulting in feed-forward Cu over-accumulation as cells grow and divide (Moreno and Docampo, 2009; Hong-Hermesdorf et al., 2014).

Here, we exploit Zn-limitation-dependent Cu over-accumulation in Chlamydomonas to understand strategies for Cu detoxification in photosynthetic organisms and to identify candidate Cu binding ligand(s) in situations of intracellular Cu overload. Survey transcriptomics and proteomics with select elemental and metabolite analysis revealed increased sulfur (S) assimilation and a remarkable 82-fold increase in cellular cysteine content, the latter being directly coupled to the accumulation of Cu. X-ray fluorescence microscopy (XFM) of Zn limited cells showed sulfur (S) enrichment in foci containing Ca and P (defining constituents of acidocalcisomes) and Cu, thus spatially linking cysteine accumulation with Cu sequestration.

## RESULTS

### Cu homeostasis is disrupted when Zn content falls below a threshold

We curated element profiles from 102 independent samples, in which wild-type Chlamydomonas cells were grown under diverse conditions (including variation in Zn and Cu nutrition) and noted that Cu over-accumulation occurred when the Zn content fell below 1 × 10^7^ Zn atoms / cell, independent of the Cu content of the medium (Figure 1a). In other words, while over-supply of Cu in the medium does not by itself result in intracellular Cu overload, Zn deficiency does. Once the cells are Zn-deficient (<1 × 10^7^ Zn/cell) the extent of Cu over-accumulation depends on how much Cu is supplied in the medium and for how long (Hong-Hermesdorf et al., 2014). Since Zn-limitation resulted in Cu overload, we tested whether Zn(II) re-supply might restore Cu homeostasis. Zn-limited cells in early exponential growth phase were collected for elemental profiling (t = −1h), following which Zn(II) was titrated into the medium (t = 0h) at Zn(II) concentrations ranging from 1 nM to 2.5 μM (the latter being the standard concentration in fresh growth medium) and the cells re-sampled for elemental profiling 24h later. In the absence of Zn re-supply, intracellular Cu content increased further, consistent with net Cu(I) uptake during the extended period of growth in Cu over-accumulating conditions. At the other extreme, supplementation with sufficient Zn(II) corresponding to a fully Zn replete medium, resulted in restoration of the normal Cu quota within a 24h period (corresponding to ~3 generations), consistent with net export of the excess Cu. As we titrate Zn(II) into the medium, we note that cells reduce Cu accumulation as a function of the Zn content of the medium (Figure 1b): at a concentration corresponding to ~ 1×10^7^ Zn /cell, there is no further net Cu accumulation (red arrow). Taken together, these data suggest that Zn-limitation below a threshold of 1 × 10^7^ per cell results in loss of function of a Zn-protein that is important for Cu homeostasis.

**Figure 1.**
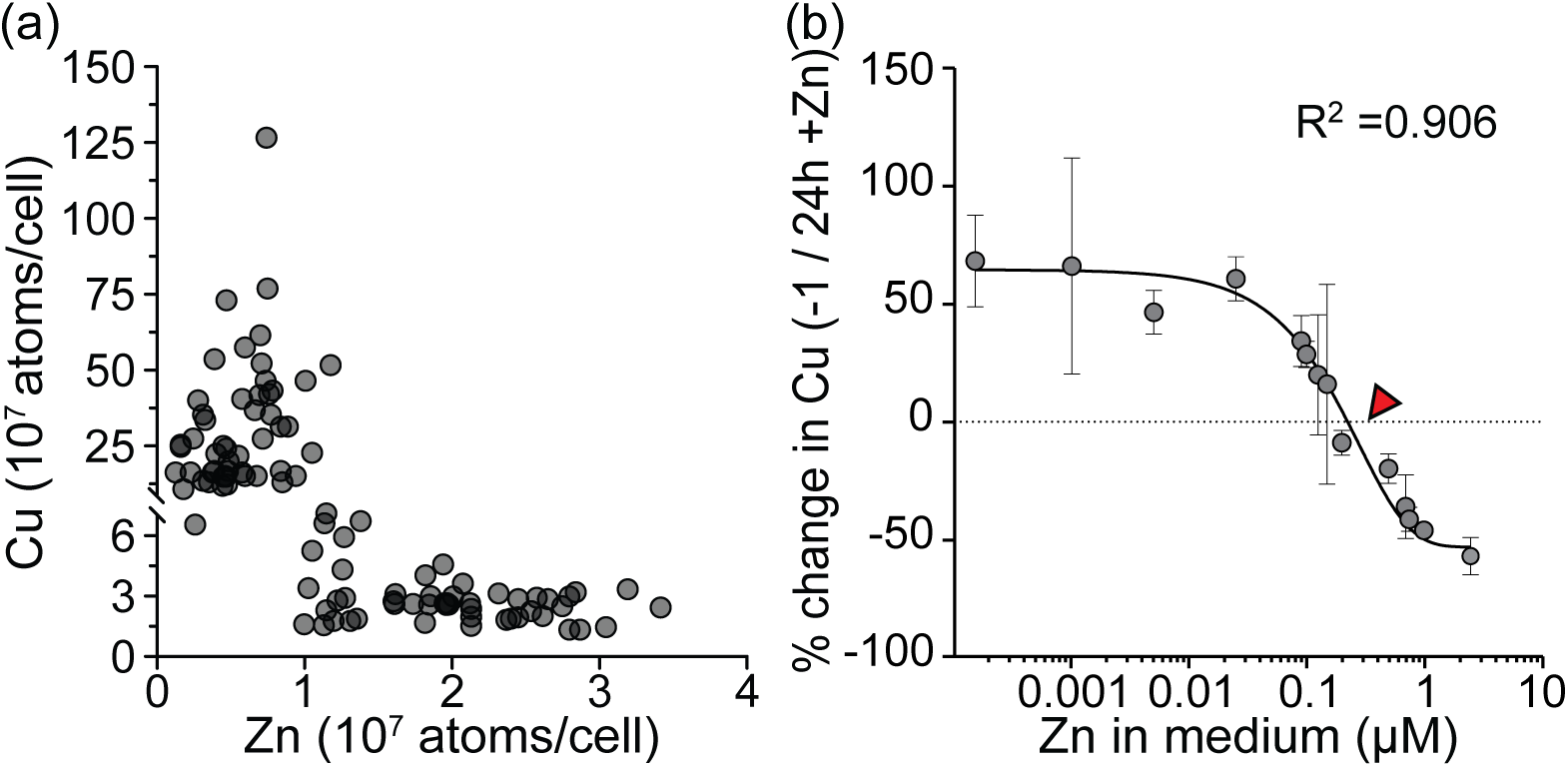
Zn deficiency induced imbalance of Chlamydomonas’ Cu content is restored at > 1 × 10^7^ Zn atoms / cell. (a) Cu and Zn content from Chlamydomonas cells was measured by ICP-MS/MS. Each circle reflects one of 102 independent samples in which Chlamydomonas cells were grown under various Zn and Cu concentrations. (b) Cells were grown in growth medium with 0 μM Zn, in the presence of replete amounts of Cu (2 μM), resulting in Zn deficiency induced Cu accumulation. After cells reached early exponential growth (1-2 × 10^6^ cells/ml), various amounts of Zn were added back to the cultures to final concentrations as indicated on the x-axis The relative change of cellular Cu content before (−1) and after (24h) Zn addback was plotted against the concentration of Zn in the medium. The amount of Cu was measured by ICP-MS/MS in three independent experiments. The white open circle corresponds to the amount of Zn in standard growth medium (2.5 μM), considered replete. The red arrow indicates the amount of Zn corresponding to 1 × 10^7^ Zn atoms / cell. The relative values with SDTEV are presented, taking the −1 sample values as 100%.

Tight regulation of Cu homeostasis is typically a balance between passive Cu uptake and active Cu export (Odermatt et al., 1992; Sancenón et al., 2003; Jung et al., 2012): over-accumulation can result from increased uptake or decreased export. To distinguish between the two possibilities, we grew Zn-limited cells in medium containing isotopically pure ^63^Cu, so that intracellular Cu was fully distinguished as ^63^Cu, and measured the Cu content (t = −1h) before cells were washed and resuspended in fresh Zn-free (-Zn) or Zn-replete (+Zn) medium containing ^65^Cu (t = 0h) as the only Cu source. Cells and spent medium were sampled after continued growth and division (24 and 72h) for measurement of ^63^Cu and ^65^Cu inside the cells and outside in the spent medium (Figure 2). The total Cu content of the Zn-limited cells increases dramatically over 3 days of growth as cells approach stationary phase, but remains low in Zn-replete cells, with only a slight increase at t=72h as cells approach stationary, as was noted before (Hui et al., 2022). The loss of intracellular ^63^Cu over time (from −1 to 24 and 72) is not different between Zn-replete vs. Zn-limited cells (Figure 2b, left panel, yellow bars). This is accompanied by gain of ^63^Cu in the spent medium over time (Figure 2b, right panel, yellow bars) and again not different between Zn-replete vs. Zn-limited conditions. This indicates that Cu export from cells is not affected by Zn nutrition. In contrast, cell-associated ^65^Cu is increased in -Zn cells relative to +Zn cells; in fact, the higher Cu content of -Zn cells is entirely attributed to uptake of ^65^Cu from the medium. The corresponding depletion of ^65^Cu from the -Zn spent medium is evident relative to the +Zn medium. These data support the existence of a balanced Cu uptake and export system in Chlamydomonas as in other organisms and identify Cu uptake as the affected target of Zn-limitation.

**Figure 2.**
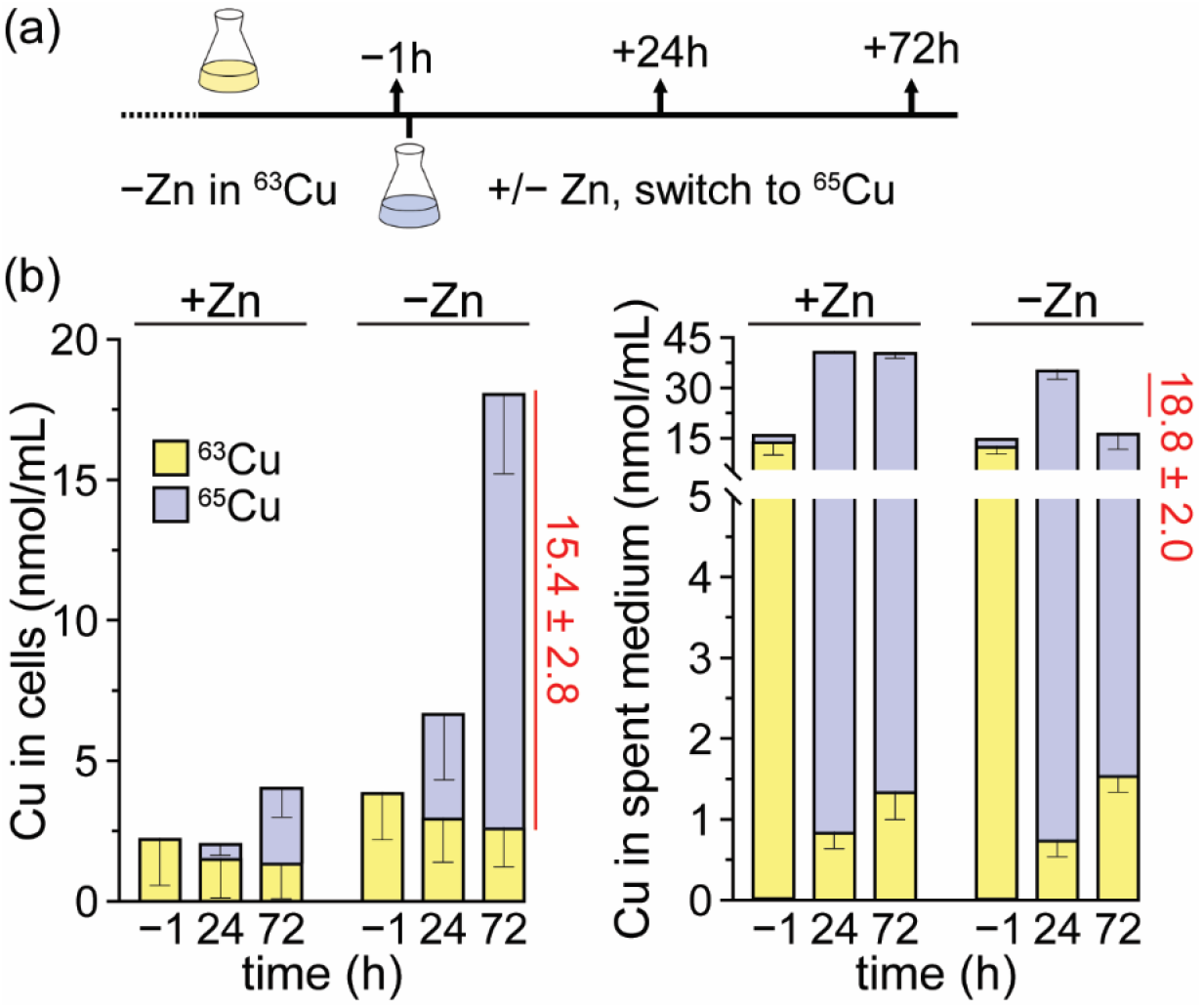
Zinc deficiency induced Cu accumulation is the result of increased Cu uptake. (a) Schematic overview of experimental design. Cells were grown in ^63^Cu containing growth medium without supplemental Zn as indicated (-Zn), collected by centrifugation and transferred to medium that was not supplemented (0 μM, −) or supplemented with Zn (2.5 μM, +). ^65^Cu was added simultaneously at this time (2 μM final concentration). (b) Cells (left) and spent media (right) were collected and isotope specific Cu content was measured by ICP-MS/MS. Shown are averages and STDEV of three independent experiments.

### A proteomic approach combined with transcriptional and elemental profiling reveals a surprising relationship between Zn limitation and S assimilation

To decipher a target of Zn-limitation that might impact Cu homeostasis, we compared transcriptomes from Zn-replete vs Zn-limited cells (Hong-Hermesdorf et al., 2014) and noted that a substantial fraction (25-40%) of the differentially accumulating transcripts under Zn-limitation were also differentially accumulated under S-limitation (Gonzalez-Ballester et al., 2008) (Figure 3a and Supplemental Table S1). Curation of the lists identified the following commonalities: decreased abundance of transcripts encoding components involved in photosynthesis, protein, DNA and RNA synthesis, and progression through the cell cycle (a not unexpected impact of nutrient limitation (Schmollinger et al., 2014)), and increased abundance of transcripts encoding components of S assimilation and S metabolism, which is not unexpected for the S limitation transcriptome, but interesting in the context of Zn limitation (Figure 3b).

**Figure 3.**
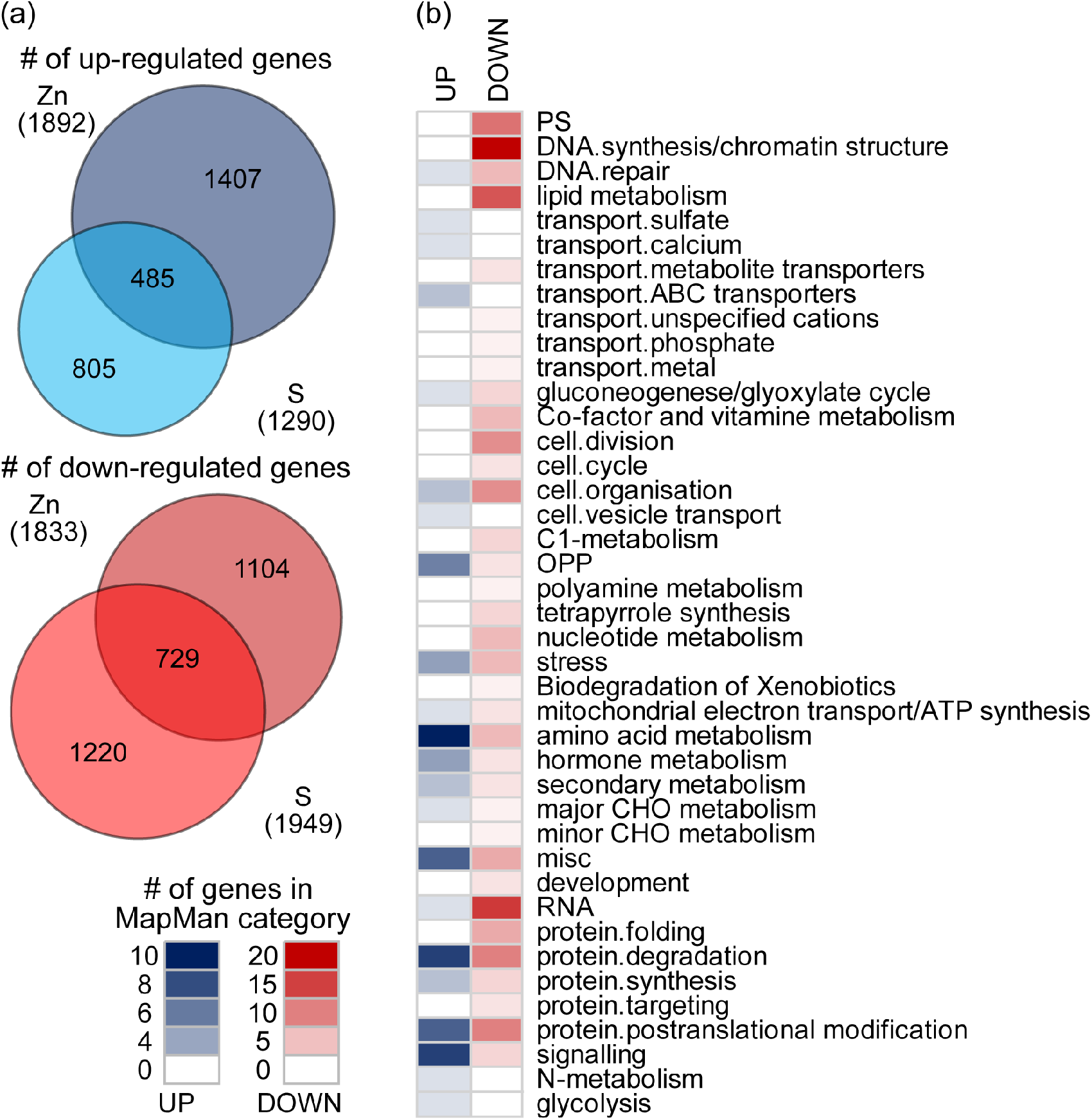
Overlapping patterns of gene expression between S and Zn limited Chlamydomonas. (a) Venn diagrams depict differentially expressed genes (DEGs) in Zn limited and S deficient Chlamydomonas. (b) MapMan annotations from genes found to be up- (blue) or down- (red) regulated in both data sets were derived using the functional annotation tool (http://pathways.mcdb.ucla.edu/algal/index.html). The number of up- and down-regulated genes in each MapMan category is displayed as a heat map. Gene expression data are derived from experiments described in Hong-Hermesdorf et al. 2014 and González-Ballester et al. 2010.

To assess whether observed changes in the transcriptome are realized at the level of the corresponding proteins, and whether they impact metabolism and physiology, we used an untargeted proteomics approach to evaluate Zn-replete vs. -limited cells. Triplicate samples of Chlamydomonas cells grown in either condition were analyzed to identify 9708 proteins, with excellent correlation between individual replicates (Supplemental Figure 1). Relative protein abundances were determined using the maximum peak intensity values (Monroe et al., 2008).

In addition, we applied the total protein ruler method (Wisniewski, 2017) to estimate absolute protein abundances in fg/cell, assuming that the total summed mass-spectrometry derived signal reflects the cumulative protein mass of the cell. This approach was previously used to determine absolute abundances of the Zn proteome in yeast (Wang et al., 2018). In our experiment, the total protein content of Chlamydomonas cells ranges between 24 pg/cell in Zn replete cultures to 16 pg/cell in Zn deplete cultures (Supplemental Figure 2a), which is in the expected range based on previous protein concentration estimates of photoheterotrophically grown Chlamydomonas (Umen and Goodenough, 2001; Hammel et al., 2018). The reduced total protein content in Zn limited cells might reflect a loss of ribosomal proteins, which are amongst the most abundant Zn containing proteins in a cell, and hence prone to degradation in Zn deficiency (Wang et al., 2018) (Supplemental Figure 2b). Besides the direct impact on the abundance of the ribosomal proteins themselves, reducing ribosome abundance has an additional impact on the capacity to synthesize new proteins, which contributes to reduced total protein content per cell. For individual proteins, absolute protein abundances of 3706 proteins were obtained using the most permissive criteria that protein quantification was supported by at least two identified peptides (Supplemental Table S2). Evaluation of these proteomic data together with the matching transcriptome data enabled inference of metabolic changes in Chlamydomonas in response to Zn deficiency.

Chlamydomonas, like many microbes, can take up inorganic sulfur and assimilate it into organic species after reduction (Pollock et al., 2005). Inducible, high affinity sulfate importers of Chlamydomonas include *SULTR2* (for <u>sul</u>fate <u>tr</u>ansporter 2)*, SLT1 and SLT2* (for <u>S</u>AC1-<u>l</u>ike <u>t</u>ransporter <u>1</u> and <u>2</u>) (Yildiz et al., 1994; Takahashi et al., 2001; Pootakham et al., 2010): all those transcripts increased in abundance by more than 100-fold in response to Zn limitation (Figure 4a, left panel). We quantified both SLT1 and SLT2 in the proteomics experiments and found that their abundances increased more than 10-fold in Zn limited cells (Figure 4a, right panel). SULTR2 peptides were not detected in these experiments. To see if the increase in S transporter abundance has an impact on S metabolism, we compared the S contents of Zn-deplete and Zn-replete Chlamydomonas cultures by elemental profiling. The analysis revealed a significant increase in intracellular S under Zn limitation, a presumed consequence of the increase in *SLT* transcripts and SLT transporters abundances in these conditions (Figure 4b). Interestingly, the increase in total cellular S content was even greater when Zn limited cells were provided with 5-fold excess of Cu (10 μM) in the medium, which further increases intracellular Cu accumulation (Figure 4b).

**Figure 4.**
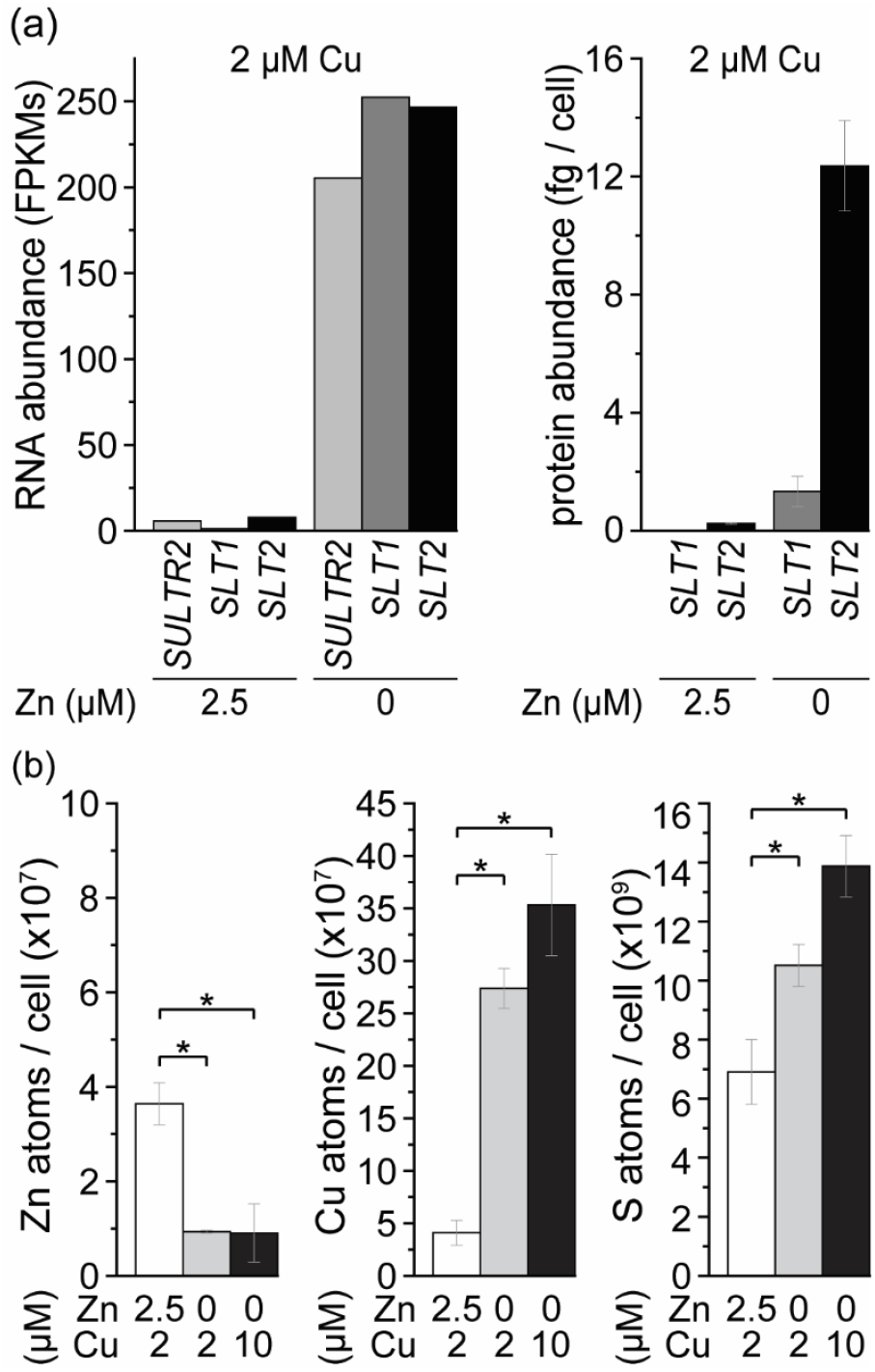
High affinity S transporters are induced in Zn limited Chlamydomonas, resulting in increased S content. (a) Shown are RNA abundance values determined by RNA-Seq in FPKMs for *SULTR2*, *SLT1* and *SLT2* and protein abundances in fg / cell for the corresponding proteins (SLT1, SLT2). Cells were grown in either Zn replete (2.5 μM) or Zn limited (0 μM) medium. Shown are averages and STDEV of three independent cultures. Significant differences were determined by one-way ANOVA followed by Holm-Sidak, p-value ≤ 0.05. (b) Wildtype cells (CC-4533) were either grown in Zn replete (2.5 μM) or in Zn limited medium (0 μM) in which 2 μM Cu (replete) or 10 μM Cu (excess) was added as indicated. Intracellular Zn, Cu and S was measured by ICP-MS/MS. Shown are averages and STDEV of three independent cultures.

We conclude that the higher S content in Zn limited Chlamydomonas is mediated by the inducible, high affinity sulfate import system and is regulated at the level of transcript accumulation. This result is reminiscent of the impact of Cd on S metabolism in yeast, where the exposure to this toxic metal draws on the intracellular pool of S metabolites owing to increased glutathione synthesis required for metal chelation (Fauchon et al., 2002).

### Cysteine as a Cu ligand - a novel, ancestral route for Cu detoxification?

Therefore, we inventoried the abundance of enzymes functioning in the biosynthesis of S-containing amino acids, glutathione, PC and its precursors, and the corresponding intermediates and metabolites in Zn-replete vs. -limited cells. Transcripts encoding enzymes for cysteine biosynthesis were increased across the board in Zn limited cells, including sulfite reductase (*SIR1*), serine *O*-acetyl transferase (*SAT1*, *SAT2*) and *O*-acetylserine (thiol)-lyase (*OASTL1*, *OASTL2*, *OASTL3*, *OASTL4*) (Figure 5a) with corresponding dramatic increases in enzyme abundances for sulfite reductase and serine *O*-acetyl transferase (Figure 5b). Absolute protein quantifications suggest that, as in land plants, *O*-acetylserine (thiol)-lyase is in vast excess (50 -100x) over serine *O*-acetyl transferase (Figure 5b); hence, it is not rate limiting for Cys production and not dramatically up-regulated like some of the other enzymes. Analysis of water-soluble metabolites using HPLC revealed remarkable increases in two pathway end-products, cysteine and y-glutamylcysteine, 82 and 10-fold, respectively, but not in the pathway intermediates (Figure 5c). On the other hand, glutathione content (measured as the sum of the reduced and oxidized forms) was decreased under Zn limitation. PC abundance was magnitudes lower than Cys abundance and did not change significantly in response to Zn limitation. Based on an absolute quantitative assessment of the intracellular Cys content using HPLC and internal standards (Figure 5c), we estimate ~ 2.82 × 10^9^ Cys molecules in a Zn limited cell.

**Figure 5.**
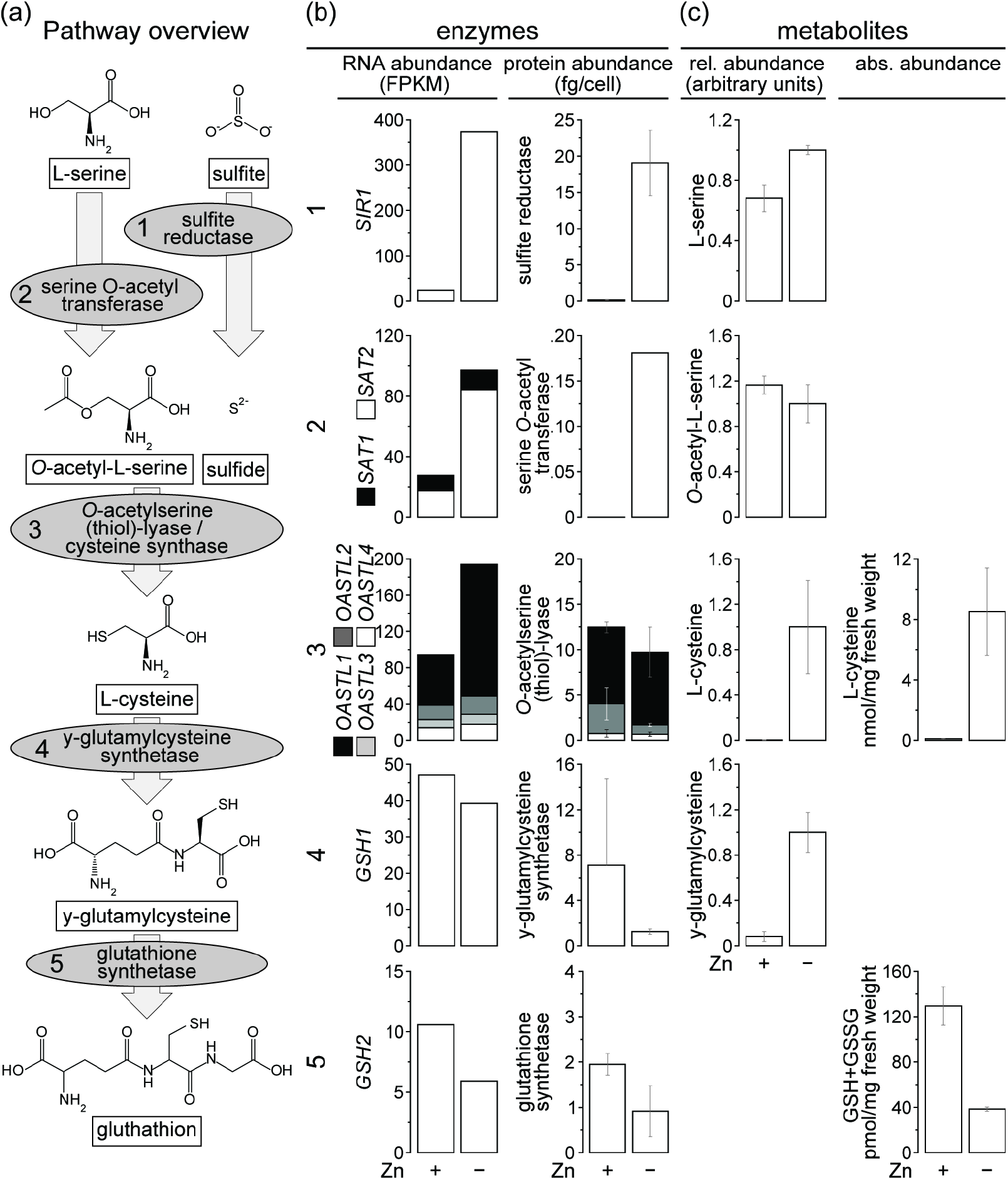
Changes in expression of enzymes involved in S metabolism result in increased Cys and γ-GluCys content. (a) Metabolites are shown in boxes and enzymes in gray ellipses. Relative (b) transcript abundances determined by RNA-sequencing (in FPKMs) and protein abundances determined by mass spectrometry (in fg/cell). Shown are averages and STDEV of three independent cultures. (c) Changes in water-soluble metabolites in Zn replete and Zn limited grown cells, analyzed by HPLC. Left row: results are shown as averages and SEM of three independent cultures, and maximum values for each end product are set to 1. Right row: absolute quantitative metabolite data based on internal standards on a per fresh weight basis. Shown are averages and STDEV of three independent cultures.

A metabolite survey of other amino acids indicates only slight or marginal increases in Zn limitation (perhaps owing to a decrease in protein synthesis), not comparable to the massive increase in Cys (Supplemental Figure 3). Interestingly, methionine (Met) levels were about 5-fold higher in Zn-limited cells (Supplemental Figure 3, Figure 6) even though methionine synthases are Zn-dependent enzymes. In the absence of vitamin B_12_ supplementation, METH1 is irrelevant, and methionine levels must depend on METE1 (Helliwell et al., 2014) whose abundance is increased by Zn-limitation, perhaps as a compensatory mechanism. Even if the enzyme is not fully metalated, the higher concentrations of precursors Cys and cystathionine might allow maintenance of Met levels or result in its accumulation. Another metabolic fate for Cys is conversion to taurine via Cys dioxygenase, whose transcripts are increased under Zn-limitation (Supplemental Figure 4), but we did not detect taurine in these analyses, consistent with its low abundance in Chlamydomonas (Tevatia et al., 2015).

**Figure 6.**
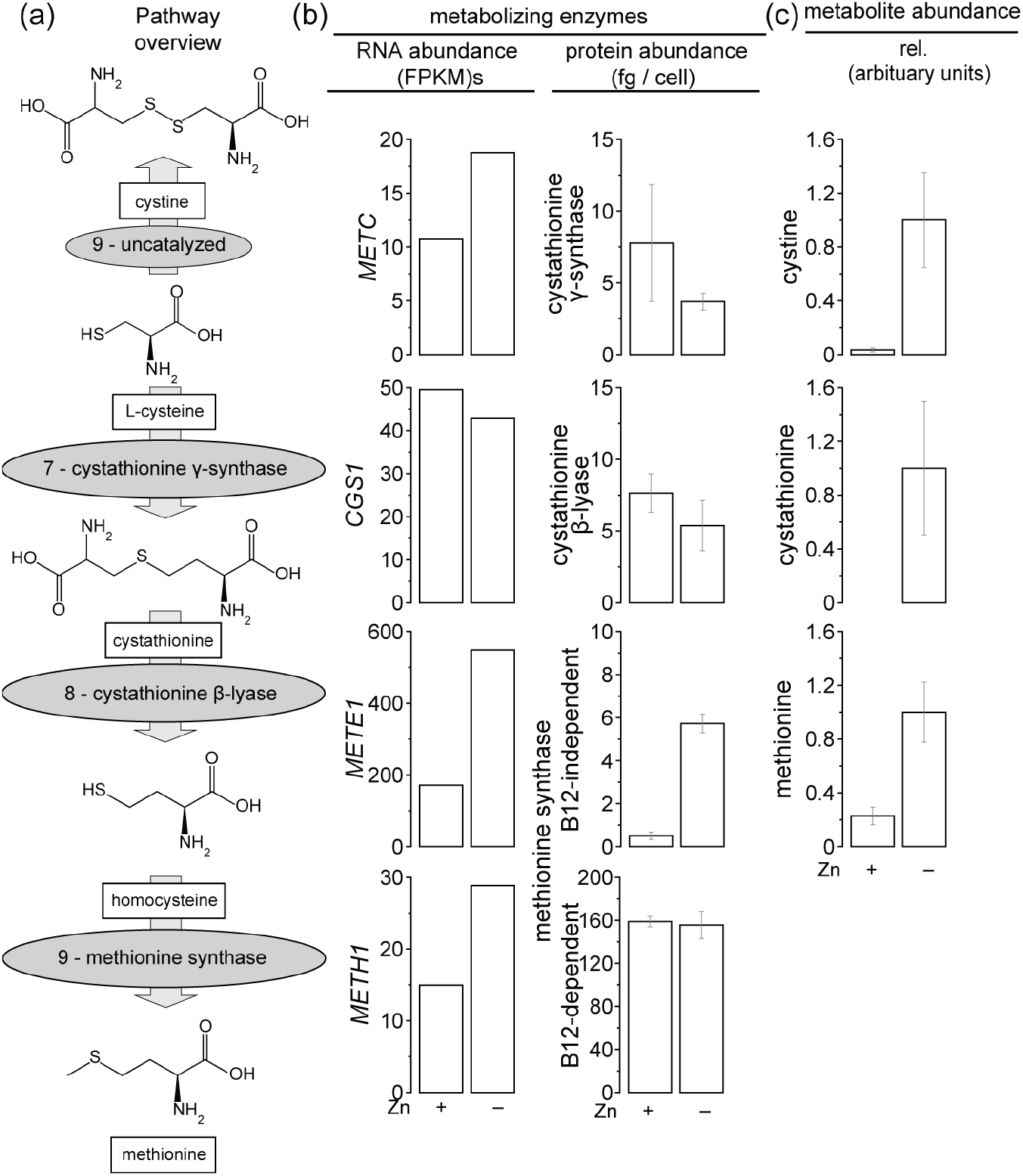
Methionine biosynthesis, which is Zn dependent, is increased in Zn limitation. (a) Metabolites are shown in boxes and enzymes in gray ellipses. (b) Relative transcript abundance determined by RNA-sequencing (in FPKMs) and protein abundances determined by mass spectrometry (in fg/cell). Shown are averages and STDEV of three independent cultures. (c) Changes in water-soluble metabolites in Zn replete and Zn limited grown cells, analyzed by HPLC. Results are shown as averages and SEM of three independent cultures, and maximum values for each end product are set to 1.

### Is Cu accumulation the consequence or the driver of cysteine accumulation?

It is possible that Cu import and cysteine biosynthesis are both mis-regulated in Zn-limited cells but otherwise are not metabolically connected. To establish a functional connection, we imposed Zn limitation on Cu deficient cells. As expected, Cu-deficient cells cannot over-accumulate Cu although they can accumulate normal amounts of Mn and Fe (Figure 7a) (Schmollinger et al.,2021). Notably, Zn/Cu co-limited cells had significantly less intracellular S as compared to Cu-accumulating cells. This phenotype was specific to S because the abundance of phosphorus (P) is unchanged between Cu-replete and Cu-deficient conditions. The observed reduction in S (2.87 × 10^9^ sulfur atoms per cell) matches closely the amount of Cys (and hence S) accumulated in a Zn limited cell (2.82 × 10^9^ cysteines per cell) (Figure 5). Most importantly, the observed reduction in S content in the Cu/Zn-limited cells is concomitant with an 88 percent reduction of cysteine as compared to the Cu accumulating cells (Figure 7b).

**Figure 7.**
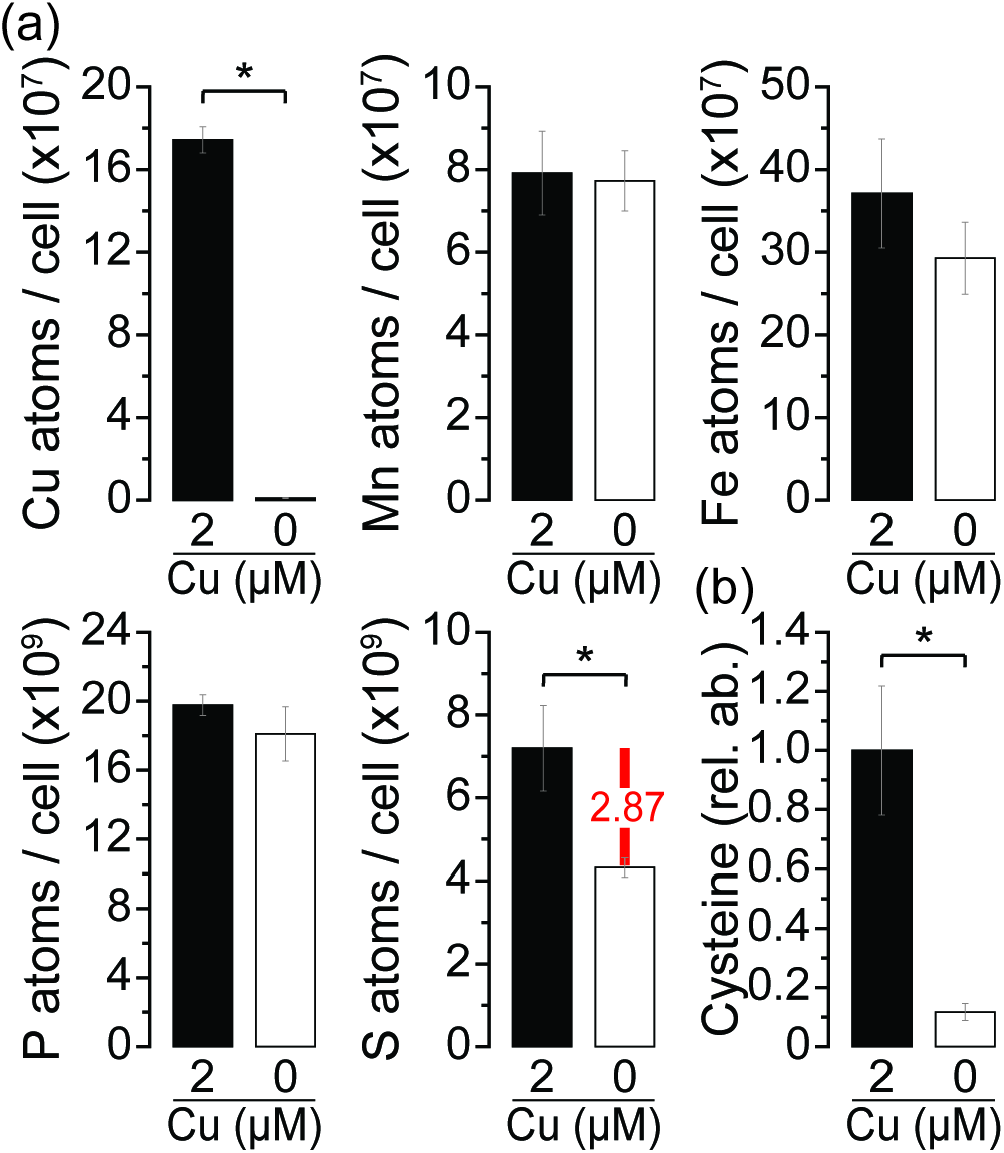
Cu is required for cysteine accumulation in Zn limited cells. Cultures were either grown in Cu deplete medium (0 μM) or Cu containing medium (2 μM) for three consecutive rounds before transfer to Zn deplete media. Cells were collected after cultures reached a density between 6 and 8 × 10^6^ cells / ml. (a) Intracellular Cu, Mn, Fe, P, and S content was measured using ICP-MS/MS. Shown are averages and STDEV of three independent cultures. (b) Relative abundance of cysteine was analyzed by HPLC. Shown are averages and STDEV of three independent replications. Significant differences were determined by one-way ANOVA followed by Holm-Sidak, p-value ≤ 0.05.

Taken together, these results functionally connect the production of high levels of cysteine with Cu accumulation, and they also underscore the induction of S uptake and cysteine biosynthesis as a very specific response to an increase in intracellular Cu content.

### S is enriched and co-localized with Cu, Ca and P inside acidocalcisomes

Previously, we had shown that the excess intracellular Cu in over-accumulating situations is sequestered into acidocalcisomes, which are lysosome related organelles containing polyphosphate (polyP) and calcium (Ca), using various elemental imaging techniques, including XFM, nanoSIMS and Cu-specific dyes (Hong-Hermesdorf et al., 2014; Schmollinger et al., 2021). Since XFM collects the fluorescence signal for a wide range of elements, not only the ones directly specified and of immediate interest, we reanalyzed previous images of Cu hyper-accumulating Zn limited cells to examine S distribution (Schmollinger et al., 2021). S is a macro-nutrient, found in all proteins as well as in the thylakoid membranes of chloroplasts, and is typically distributed throughout the cell. When we compared Zn-limited (0 Zn) vs. Zn-replete (2.5 μM Zn) cells, we identified distinct foci of S concentration in the former but not latter (Figure 8a). More importantly, these S foci are co-localized with Cu, consistent with complexation of the sequestered Cu by a S-containing molecule. As previously noted, Cu and S are co-localized with Ca (Figure 8b) and P (Supplemental Figure 5), indicating that the excess S in Zn-limited cells is also in the acidocalcisome. These data spatially connect the accumulation of Cu and S and our results are consistent with prior EXAFS analyses that indicated S and N ligands for the sequestered Cu(I) (Hong-Hermesdorf et al., 2014).

**Figure 8.**
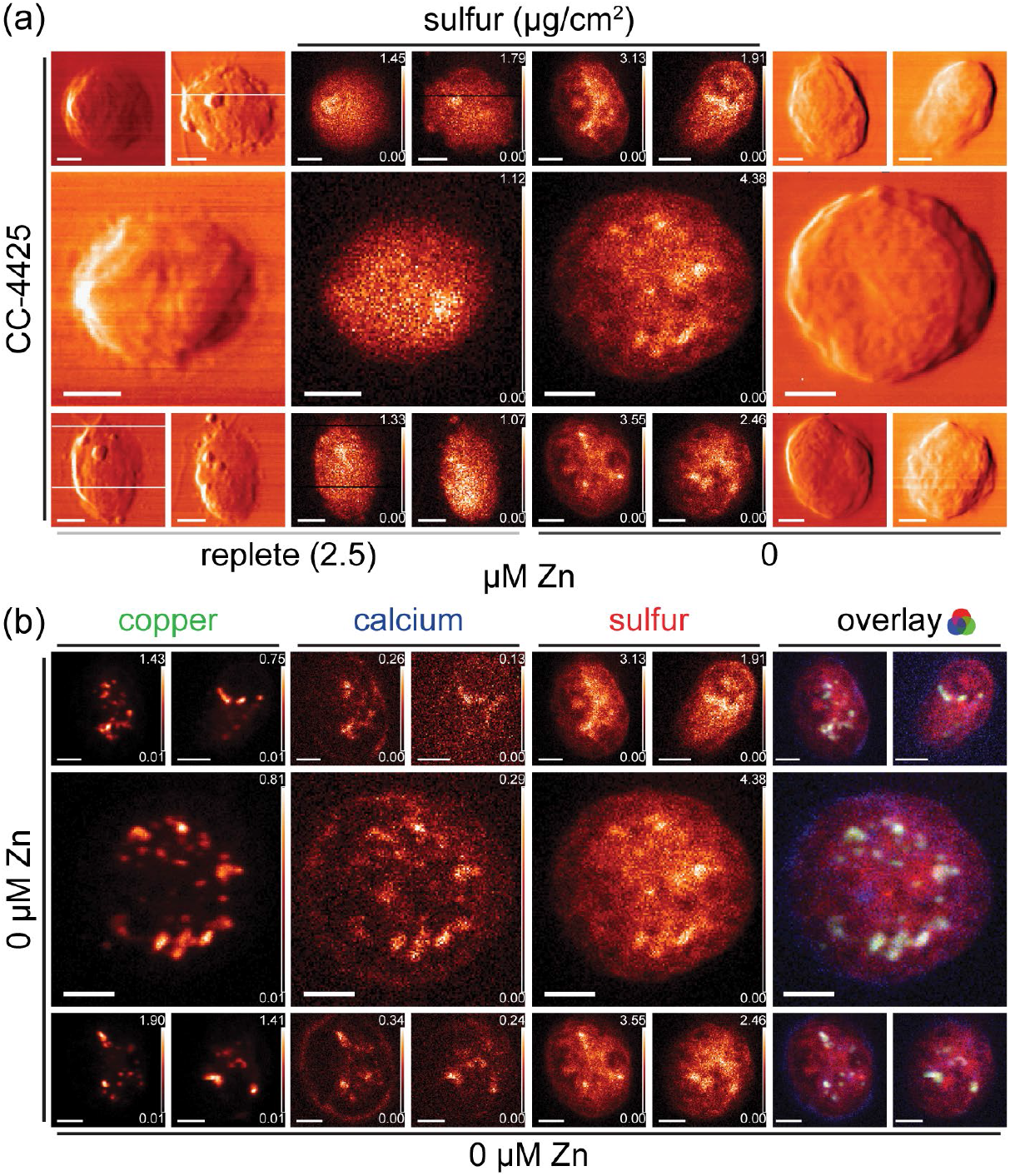
Cu co-localizes with P, Ca and Sulfur. Shown are X-ray differential phase contrast and false color 2 D elemental maps for Ca (blue), S (green), P (red) as well as an overlay for all three elements (merge). Minimum and maximum area concentration are displayed above each element. Cells were grown in 2.5 or 0 μM Zn and fixed on X-ray sample supports were raster scanned at the APS Bionanoprobe. X-ray fluorescence data were measured in flyscan mode using an energy dispersive detector, fluorescence spectra fitted at each point of the scan, and the elemental area concentration determined from measuring X-ray standards. The scan area was 13 × 12 um; pixel size: 70 nm; dwell time: 200 ms.

## DISCUSSION

### Compartmentation and chelation

In undisturbed cells, the Cu quota is predominantly determined by the abundance of the major copper sinks, which are its most abundant cuproproteins. In Chlamydomonas, as in other photosynthetic organisms, over 90% of intracellular copper is found associated with only two proteins, mitochondrial cytochrome *c* oxidase and plastid-localized plastocyanin (Kropat et al.,2015). When Cu homeostasis is disrupted, as in Zn deficient cells, leading to an up to 40-fold increase in Cu content, most of the intracellular Cu is no longer protein-associated. Instead, that Cu is potentially present in the cell as readily exchangeable species, which can be deleterious. Besides the problem of redox reactivity, there is the problem of mis-metalation because cuprous and cupric ions are competitive with Zn for metal binding sites in proteins, especially when the Zn concentrations are low as in a Zn-deficient cell (Imlay, 2014). A solution to this problem is Cu sequestration via compartmentation and / or chelation by analogy to phytochelatin-based heavy metal detoxification in land plants and animals (Vatamaniuk et al., 2001; Sharma et al., 2016). In this work, we used systems level analysis of mis-regulated Cu homeostasis to identify candidate Cu(I)-binding ligands and to further probe the biochemistry of Cu(I) storage sites in Chlamydomonas, Cys and the acidocalcisomes, respectively.

When reactive metal species are sequestered within a membrane-bound organelle, they are physically separated from enzymes with which they may associate adventitiously (a real problem for Cu at the top of the Irving-Williams series) and from labile metabolites or macromolecules (especially in the presence of oxygen) (Imlay, 2014). In Zn deficiency in Chlamydomonas, Cu(I) sequestration takes place in the acidocalcisomes (Malasarn et al., 2013; Hong-Hermesdorf et al.,2014; Schmollinger et al., 2021), which are acidic, lysosome-related organelles (Huang et al.,2011). We extended our previous work by assessing whether the defining chemistry of the acidocalcisome, namely high polyP and Ca contents, is required for Cu over-accumulation as they are indeed required for Mg and Mn accumulation in yeast and Chlamydomonas (Klompmaker et al., 2017; Tsednee et al., 2019). The Chlamydomonas VTC complex is required for polyP synthesis and Ca accumulation (Aksoy et al., 2014; Gerasimaitė et al., 2014). When we tested a *vtc1* mutant (defective in polyP synthesis), we noted that, as in yeast, the P and Ca contents are drastically reduced, but the high Cu accumulation in Zn-limited cells persists (Supplemental Figure 6) and (Schmollinger et al., 2021). Therefore, we conclude that the eponymous acidocalcisome biochemistry is not required for Cu over-accumulation in Chlamydomonas. It must rather be driven by chelation.

### Cysteine: a novel Cu(I) ligand?

A significant rise in internal S levels, the induction of S transporters and EXAFS data (Hong-Hermesdorf et al., 2014) in Zn limited Chlamydomonas cells points towards a S-containing ligand in Cu detoxification. Although PCs, previously shown to function in heavy metal detoxification in Chlamydomonas, are an obvious candidate, no increase in GSH or PC abundance was observed in Zn-deficient, Cu-over-accumulating cells (Figure 5). Instead, the amino acid Cys increased >80 fold in the cell with a corresponding increase in the total S content (Figure 4, 5). Transcriptome and proteome analyses are consistent with this observation: increased expression of sulfate assimilation (SLTs, SULTR2), reduction (SIR1) and S incorporation into Cys (SAT1) (Figure 5), but no change in the expression of phytochelatin synthesis enzymes.

The increase in S content is strikingly similar to the increase in Cys content and is not determined solely by the absence of Zn, but rather dependent on Cu availability and driven by Cu supply (Figure 7). If we assume that cysteine detoxifies Cu in a Zn limited cell, and based on previous EXAFS data suggesting a mononuclear Cu species with 1S and 2 O/N coordinating ligands (Hong-Hermesdorf et al., 2014), we would expect a 1:1 stoichiometry between Cu(I) and cysteine. Instead, we estimate a 10-fold excess of cysteine as compared to the amounts of Cu. We can explain this discrepancy by invoking higher Cys concentrations as a thermodynamic driving force for stable Cu(I)-Cys association, which is necessary because of the relatively low binding affinity between Cu(I) and Cys (Walsh and Ahner, 2013).

We suggest that ancient metal detoxification mechanisms may have relied on direct binding of metal ions by amino acids or amino-acid-derived metabolites. Indeed, Cys is known to be compartmentalized in the lysosome in mammalian cells and excess Cys in yeast perturbs Fe bioavailability, documenting the connection between amino acids (especially Cys) and redox active transition metals (Abu-Remaileh et al., 2017; Hughes et al., 2020). Proline is another amino acid implicated in metal, especially Cd, stress tolerance in plants and algae (Siripornadulsil et al.,2002; Bačkor et al., 2004). Compounds derived from amino acids, like γ-glutamylcysteine (γ-GC), PCs and ergothioneine, may represent more recent additions to the metal chelating repertoire of cells. Ergothioneine, derived from histidine, is a competent Cu chelator found in select filamentous fungi (Genghof, 1970; Zhang et al., 2022) and some cyanobacteria (Genghof, 1970; Pfeiffer et al., 2011; Zhang et al., 2022). In yeast, the dipeptide of γ-glutamylcysteine (γ-GC), formerly also known as cadystins (Kondo et al., 1985; Hayashi et al., 1991)which also accumulates in Zn limited Chlamydomonas (likely driven by the high Cys content), but to a much lower level than Cys, can also chelate Cu. The more complex polyvalent ligands like PCs are likely newer innovations, selected for higher affinity metal binding. In Chlamydomonas, glutathione (GSH) is adequate for detoxifying Hg(II) ions because of the strength of the Hg-S interaction, but Cd(II) detoxification requires polyvalent PCs (Howe and Merchant, 1992; Hu et al., 2001; Suárez et al.,2010). Induction of PC synthesis in Chlamydomonas species occurs concomitant with an increase in cysteine but that increase is order of magnitudes lower than what we observed in this work.

### Regulation of S metabolism

The Chlamydomonas S regulon is well-described (Sanz-Luque and Grossman, 2023). A first line of response is up-regulation of the plant-type *SLTs* and the animal type *SULTR2* for sulfate assimilation, *APR*, *SIR* and *SATs* for sulfate reduction and incorporation into organic species. followed by a “second tier” set of genes that include *ARS1*, *ARS2*, *ECPs* and *LHCBM9* transporters, perhaps in response to more severe S-deficiency. The separation of output, speaking to a branched signaling pathway, is recapitulated in Zn-deficiency where the focus is on the immediate early S-deficiency response and the enzymes of Cys biosynthesis (Figure 3 and Supplemental Figure 7).

Sulfate assimilation and cysteine synthesis are regulated by S accessibility in the green alga Chlamydomonas, and algae that experience a lack of S in the growth medium induce various S assimilation pathways (Yildiz et al., 1994). One key regulator of the S-deficiency response is *SAC1* as *sac1* mutants exhibit reduced levels of sulfate uptake and are unable to synthesize, for example, the two periplasmic aryl sulfatases (Davies et al., 1996; Takahashi et al., 2001; Ravina et al., 2002; Zhang et al., 2004; Gonzalez-Ballester et al., 2008). It is surprising that part of this genetic program is also reminiscent in Zn deficient cells even though these cells experience an excess supply of external S. Our results suggest that two complementary signaling mechanisms might have evolved that either sense intra- or extracellular S deficiency. The first signaling pathway is very well-studied: once external S bioavailability is limited, cells are responding by up-regulating different aspects of S assimilation, including an increase in expression of high affinity sulfate uptake pathways and of, for instance, aryl sulfatases which mobilize alternative S sources. The situation in Zn limited grown Chlamydomonas is different: external S supply is not limited, but internal S demanding pathways are strained because of the draw of internal S to Cys. Cys is a known regulator of S metabolism and if its abundance is strongly reduced, for example because of speciation, then S limitation is signaled. The difference between both mechanisms becomes apparent by the lack of induction of genes encoding aryl sulfatases in Zn limited vs S limited cells (Supplemental Figure 7). Somehow the cells sense an increased, intracellular demand for S, while not seeing the need to induce S assimilation pathways that allow to mobilize alternative S sources (as external S is in surplus supply).

### Cu and Zn connection

The molecular connection between Cu and Zn metabolism is physiologically relevant because *crr1* mutants (that are unable to activate the Cu-deficiency regulon) cannot grow in Zn-deficient medium (Malasarn et al., 2013). A connection is also evident in cyanobacteria where the absence of a Cu chaperone increases Zn bioavailability (Dainty et al., 2010), and in humans where Zn overload results in Cu-deficiency (Bremner and Beattie, 1995). The mechanistic details underlying these connections are not yet completely elucidated; in this work, we suggest a connection through cysteine and sulfur metabolism.

## ACKNOWLEDGMENTS

This work was supported by a grant to SSM from the National Institutes of Health (GM42143). Proteomics was performed at the Environmental Molecular Sciences Laboratory (Ringgold ID 130367), proposal ID 49840, a Department of Energy Office of Science User Facility at Pacific Northwest National Laboratory (PNNL) in Richland, WA, sponsored by the Office of Biological and Environmental Research. Trent R. Northen (for amino acid profiles) was supported by the U.S. Department of Energy, Office of Science, Office of Biological and Environmental Research (DE-SC0018301). X-ray fluorescence microscopy used resources of the Advanced Photon Source, a U.S. Department of Energy (DOE) Office of Science user facility operated for the DOE Office of Science by Argonne National Laboratory under Contract No. DE-AC02-06CH11357. Stephan Clemens (for Cys and Cys metabolite quantification) gratefully acknowledges financial support by the Deutsche Forschungsgemeinschaft (CL 152/7-2).

## AUTHOR CONTRIBUTIONS

D.S., S.S. and S.S.M designed the research; D.S., S.R.S., K.H., H.W.L., Y.H., C.H., S.O.P., C.D.N., S.Ch. performed research; S.O.P., S.Cl., S.Ch., T.R.N., M.S.L. contributed new analytic/computational tools; D.S., K.H., S.S.M., S.S., S.O.P., HWL analyzed data; D.S. and S.S.M. wrote the paper.

## MATERIAL AND METHODS

### Strains and culture conditions

*C. reinhardtii* strains CC-4533 and CC-4532, were cultured in Tris acetate-phosphate (TAP) medium with revised trace elements (Kropat et al., 2011). Cultures were incubated at 24 °C with constant agitation (160 rpm) in an Innova incubator (New Brunswick Scientific, Edison, NJ) in continuous light (~100 μmol of photons per m-2·s-1) provided by cool white fluorescent bulbs (4100 K) and warm white fluorescent bulbs (3000 K) in a ratio of 2:1.

### Elemental content measurements by ICP-MS/MS

1 × 10^8^ cells from all cultures were collected by centrifugation at 3,500 × g for 3 min and washed three times with 1 mM Na2-EDTA, pH 8.0, to remove metals associated with the cell surface and twice with Milli-Q water. After removing the remaining water by brief centrifugation, cell pellets were digested with 70% nitric acid at room temperature overnight and at 65 °C for 2 h. Digested samples were diluted with Milli-Q water to a final nitric acid concentration of 2% (v/v). To measure metal content of culture medium, aliquots of the medium were treated with nitric acid and diluted with Milli-Q water to reach a final concentration of 2% nitric acid (v/v). Elemental analysis was measured by inductively coupled plasma mass spectrometry (ICP-MS/MS) on an Agilent 8800 or 8900 Triple Quadrupole instrument, using three standards for calibration (an environmental calibration standard (Agilent 5183–4688), phosphorus standard (Inorganic Ventures CGP1), and sulfur standard (Inorganic Ventures CGS1)) and two internal standards (^89^Y and ^45^Sc, Inorganic Ventures MSY-100PPM and MSSC-100PPM, respectively). Elements were determined in MS/MS mode and measured in a collision reaction cell using helium for the measurement of ^55^Mn, ^63^Cu, and ^66^Zn; hydrogen for ^56^Fe and ^40^Ca; and oxygen for ^31^P and ^32^S as the cell gas, as described previously (69). Each sample was measured in four technical replicates, and variation between the technical replicates did not exceed 5%.

### Protein digestion and peptide detection by LC-MS/MS

We collected exactly 10^8^ cells of Chlamydomonas strain CC4533 by centrifugation at 1,450 g at 4°C for 4 min. The cell pellet was washed once with 1 ml 10 mM Na-Phosphate buffer, pH 7.0, resuspended in 200 μl 10 mM Na-Phosphate buffer, pH 7.0, flash frozen in liquid nitrogen and stored in −80° until further processing. Samples were thawed and Urea was added to a final concentration of 8 M urea and the protein lysate was transferred to 2mL snap-cap centrifuge tubes (Eppendorf, Hamburg, Germany) with 0.1 mm zirconia beads and bead beat in a Bullet Blender (Next Advance, Averill Park, NY) at speed 8 for 3 minutes at 4°C. After bead beating, the lysate was spun into a 15mL Falcon tube at 2000 g for 10 minutes at 4°C. The supernatant was removed to a clean tube. Protein concentration was determined by BCA assay (Thermo Scientific, MA, USA). Urea and dithiothreitol were added to all samples at a final concentration of 8 M and 10 mM, respectively before incubation at 60°C for 30 min with constant shaking (800 rpm). All samples were then diluted 8-fold with 100 mM NH_4_HCO_3_ and 1 mM CaCl_2_ and digested with sequencing-grade modified porcine trypsin (Promega, WI, USA) provided at a 1: 50 [w/w] trypsin-to protein ratio for 3 h at 37°C. Digested samples were desalted using a 4-probe positive pressure Gilson GX-274 ASPEC™ system (Gilson Inc., WI, USA) with Discovery C18 100 mg/mL solid phase extraction tubes (Supelco, MO, USA) as follows: columns were pre-conditioned with 3 ml methanol, followed by 2 ml 0.1%trifluoroacetic acid (TFA) in water. Samples were then loaded onto 3 columns, followed by 4 ml 95:5 water: acetonitrile (ACN) 0.1% TFA. Samples were eluted with 1 ml 20:80 water:ACN 0.1% TFA, and concentrated to a final volume of ~100 μl in a Speed Vac. After determination of peptide concentration by BCA assay, samples were diluted to 0.25 μg/μl with nanopore water for LCMS/MS analysis (LC part: LC column of fused silica [360 μm x 70 cm] handpacked with Phenomenex Jupiter derivatized silica beads of 3 μm pore size (Phenomenex, CA, USA); HPLC part: HPLC NanoAcquity UPLC system (Waters, MA, USA); MS part: Q Exactive Plus mass spectrometer (Thermo Fisher, MA, USA)). Samples were loaded onto LC columns with 0.05% formic acid in water and eluted in 0.05% formic acid in ACN over 100 min. Twelve high resolution (17.5K nominal resolution) data dependent MS/MS scans were recorded in centroid mode for each survey MS scan (35K nominal resolution) using normalized collision energy of 30, isolation width of 2.0 m/z, and rolling exclusion window lasting 30 seconds before previously fragmented signals are eligible for re-analysis. Unassigned charge and singly charged precursor ions were ignored.

### MS/MS data search

The MS/MS spectra from all LC-MS/MS datasets were converted to ASCII text (.dta format) using MSConvert (http://proteowizard.sourceforge.net/tools/msconvert.html) which precisely assigns the charge and parent mass values to an MS/MS spectrum as well as converting them to centroid. The data files were then interrogated via target-decoy approach (Elias and Gygi, 2010) using MSGFPlus (Kim and Pevzner, 2014)using a +/- 20 ppm parent mass tolerance, partial tryptic enzyme settings, and a variable posttranslational modification of oxidized Methionine. All MS/MS search results for each dataset were collated into tab separated ASCII text files listing the best scoring identification for each spectrum. Max intensity peak values (MASIC values) were calculated as summed mass of peptides for a respective protein. Protein abundances (in fg /cell) were calculated using the total protein mass of each samples.

### Relative quantitative amino acid and metabolite extraction and analysis

Exactly 5×10^7^ cells of Chlamydomonas wildtype strain CC4533 were collected by centrifugation for 4 min, 4°C and 1650 *g*. Cell pellets were flash frozen in liquid nitrogen and stored at −80°C until processed. *Chlamydomonas reinhardtii* cell pellets were lyophilized and bead beat for 15 s to break the cell walls. A volume of 1 mL of methanol was added to the cell pellets followed by sonicating in a bath sonicator for 10 mins. Samples were centrifuged at 9000 g for 3 mins to remove any insoluble residue. The resulting supernatants were evaporated under vacuum at room temperature until dry and then redissolved in 150 μl methanol containing internal standards. The solution was filtered through 0.22 μM centrifugal membranes (Nanosep MF, Pall Corporation, NY, USA) and subsequently analyzed using liquid chromatography-mass spectrometry (LC-MS). Samples were analyzed using an Aglient 1290 series HPLC installed with a HILIC-Z column (150 mm ×2.1 mm, 2.7 μm, 120 Å, Agilent Technologies, Santa Clara, CA, USA). The aqueous and organic mobile phases were LC-MS grade water with 5 mM ammonium acetate and 0.2% acetic acid (A) and 95% acetonitrile with 5 mM ammonium acetate and 0.2% acetic acid (B). The column was equilibrated with 100% B for 1 min, followed by a linear gradient to 89% B by 11 min, 70% B by 15.75 min, and 20% B by 16.25 min. The mobile phase was held for 2.25 min and back to 100% B in 0.1 min. The flow rate was set to 0.45 mL/min. MS data were acquired using a Q Exactive mass spectrometer (Thermo Fisher Scientific, San Jose, CA, USA) in positive mode. MS/MS spectra were collected using Q Exactive data dependent Top2 method at 17,500 resolution using stepped collision energies of 10, 20 and 40 eV. Metabolic data were analyzed using Metatlas toolbox to obtain the extracted ion chromatography and peak area (Yao et al.,2015). Identification was performed by comparing MS/MS fragments and retention time with an in-house library of >3000 authentic standards. At least two orthogonal identification criteria including accurate mass (<5 ppm), retention time, MS/MS fragments were used for metabolite identification as provided in the Supplementary Table. Internal standards were used to ensure the retention time was consistent among samples. All the raw and processed data have been deposited in the Joint Genome Institute’s Genome Portal (Project ID: 1281190).

### Targeted, quantitative thiol analysis of cysteine and phytochelatins

Approximately 10^8^ cells were collected by centrifugation for 5 min at 4°C and 1650 g. Fresh weights were collected for each sample before flash freezing in liquid nitrogen and further processing. Thiol measurements were done as described previously in (Minocha et al., 2008). After lyophilization, 3 μl extraction buffer (0.1% v/v trifluoroacetic acid (TFA), 6.3 mM diethylenetriaminepentaacetic acid (DTPA), 40 μM N-acetylcysteine (NAC) as internal standard) per mg lyophilized cell pellet were added and vortexed for 1 min. For analyses targeting cysteine, 60 μl extraction buffer were added per mg cell pellet. Samples were incubated on ice for 15 min and vortexed for 5 seconds every 5 min. Following centrifugation (13000 rpm at 4°C for 15 min), 31.25 μl supernatant was transferred to brown tubes. 77 μl solution A (200 mM EPPS, 6.3 mM DTPA, pH 8.2) and 3.125 μl solution B (20 mM Tris-(2-carboxyethyl)-phosphine (TCEP) dissolved in 200 mM EPPS, pH 8.2) were added and incubated for 10 min at 45 °C. For samples w/o TCEP, solution B was replaced by 3.125 μl 200 mM EPPS, pH 8.2. Derivatization was started by adding 2.5 μl 50 mM monobrombimane (dissolved in acetonitrile). After 30 min at 45 °C, derivatization was stopped by the addition of 12.5 μl 1 M methanesulfonic acid (MSA). HPLC conditions were described as in (Kühnlenz et al., 2016). Derivatized thiols were separated on a reprosil 100 C18 column (250 x 4.6 mm, 5 μm particle size) at 40 °C with an acetonitrile gradient. Thiols were detected using a fluorescence detector (FP 2020, JASCO Germany GmbH) set at excitation and emission wavelengths of 380 and 470 nm, respectively. The JASCO ChromPass Chromatography Data System (JASCO Germany GmbH, version 1.8.6.1) was used to analyze the data. Thiol quantification was performed based on authentic standards after normalization to the internal NAC standard.

### X-ray fluorescence microscopy (XFM)

3×10^6^ Chlamydomonas cells of a culture during logarithmic growth phase (at a cell density between 1-5×10^6^ cells/mL) were collected by centrifugation at 16,100 *g* at room temperature for 15 sec in a 1.5-mL Eppendorf tube. The cell pellet was washed twice in 1 x phosphate-buffered saline (PBS) buffer (137 mM NaCl, 2.7 mM KCl, 10 mM Na_2_HPO_4_, 1.8 mM KH_2_PO_4_) and resuspended either in 0.3 mL Milli-Q water for vitrification or 4 % para-formaldehyde (EM grade #15710 from EMS) in 1x PBS for chemical fixation. 100 μL of the cell suspension was then applied to a poly-L-Lysine coated silicon nitride membrane window (5 × 5 × 0.2 mm frame, 2 × 2 × 0.0005 mm Si3N4 membrane, Silson) and allowed to settle for 30 min at room temperature. After the supernatant was removed by gentle suction, the window was either directly mounted into an FEI Vitrobot Mark IV freezer for vitrification in liquid ethane (blot time 3 sec, blot force 2, blot total 1, wait and drain time 0 sec) or washed twice with 1x PBS, fresh 0.1 M Ammonium Acetate and Milli-Q water. Windows with chemically fixed cells were air dried and stored at room temperature, windows with vitrified cells were kept in liquid nitrogen storage until measurement at low temperatures. The data presented here are from four individual sessions (48-72 h measurement time each) at the Bio nanoprobe (Chen et al., 2014), over the period of two years, during which the analyzer was localized either at beamline 21-ID-D or beamline 9-ID-B at the Advanced Photon Source (APS) of the Argonne National Laboratory. Direct excitation of atomic K transitions was aimed for by tuning the incident X-ray energy to 10 keV, which allows efficient excitation of elements in the elemental table up to Zn (z=30). Individual, large Chlamydomonas cells can be up to 15 μm in diameter before division, which is well below the 50-100 μm that can be penetrated by XFM with high spatial resolution (Wagner et al., 2005). Whole cells were therefore analyzed without sectioning. Two rounds of coarse scans were performed to identify cell locations on the Si3N4 windows, with a pixel size between 0.5-2 μm in x and y direction. Then, a high-resolution scan of the individual cells was initiated for all images; a spatial resolution between of ~80 nm with a dwell time of 200 – 300 ms per pixel was used. With an average scan area of 10 x 10 μm, this results in measurement times up to 3 h per cell for each high resolution. By normalizing the recorded fluorescent spectrum with data obtained from a calibration standard thin film (AXO DRESDEN GmbH), fully quantitative maps are obtained showing the distribution for each of the individual elements.

