## Supplemental Figures and legends for "Cysteine: an ancestral Cu binding ligand in green algae?"

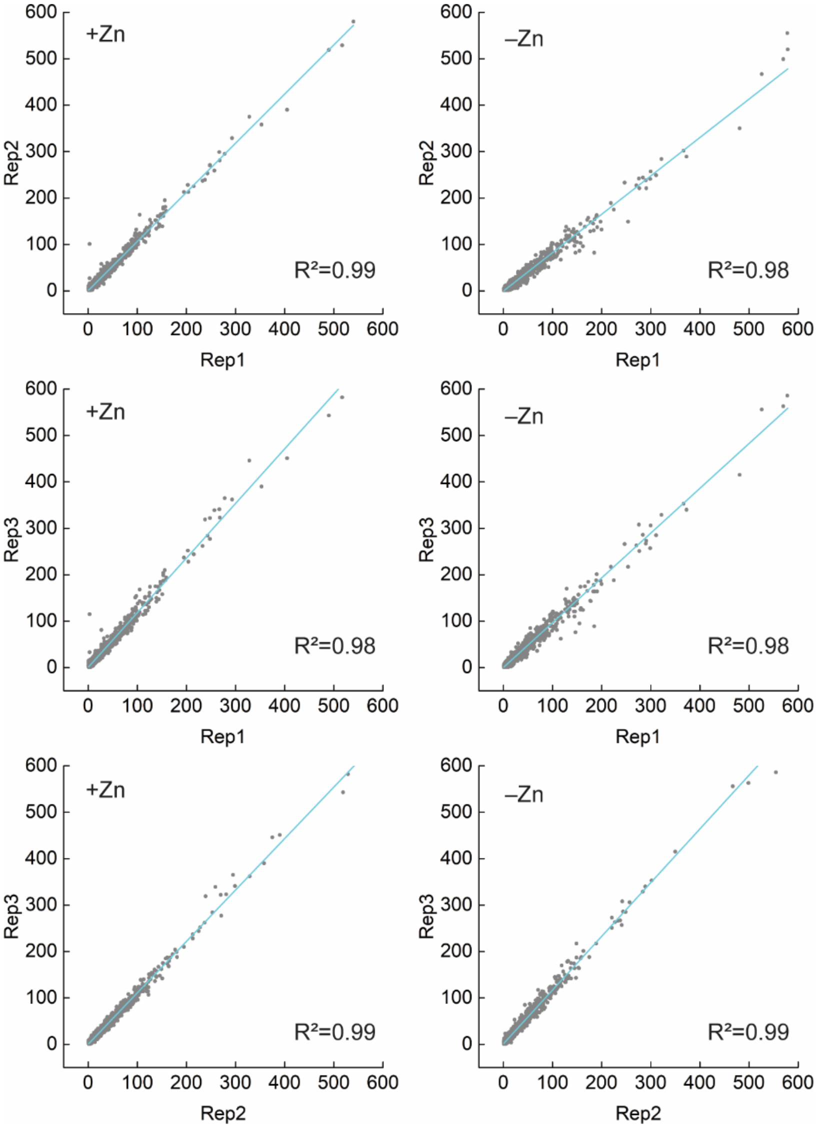


**Supplemental Figure 1. Quantitative recovery of proteins by untargeted proteomics.** Assessment of reproducibility based on spectral counts detected for all proteins between replicates and conditions. Cyan line: linear regression fit.

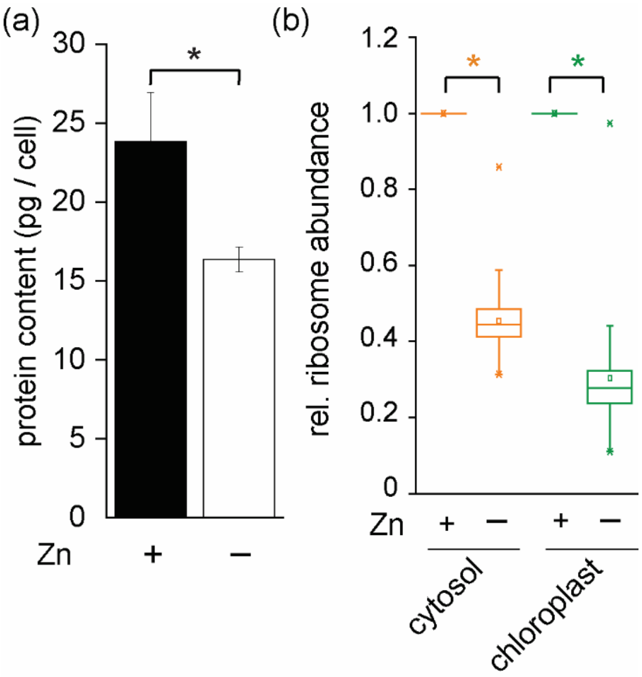


**Supplemental Figure 2. Total cellular protein content is decreased in zinc deplete grown Chlamydomonas cells.** Wildtype cells (CC4533) were either grown in Zn replete media (2.5 µM Zn, +) or in Zn deplete media (0 µM Zn, –) as indicated. (a) Cell counts were determined using a hemocytometer and protein concentrations were estimated using a BCA assay. Shown are averages and STDEV of three to four independent replications. (b) relative protein abundances of cytosolic (orange) and chloroplastic (green) localized ribosomes. Significant differences were determined by one-way ANOVA followed by Holm-Sidak, p-value ≤ 0.05.

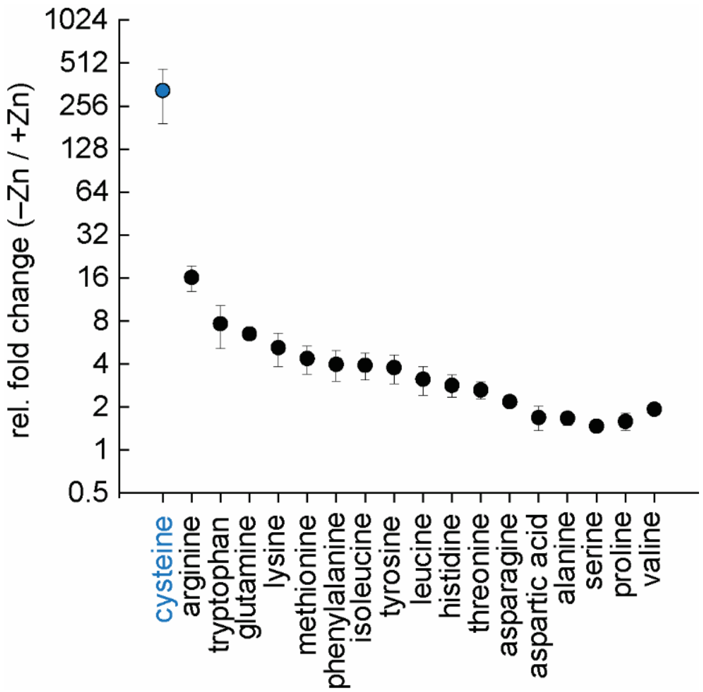


**Supplemental Figure 3. The change in abundance in L-Cysteine in response to Zn limitation is unique.** Fold induction of amino acids as a function of Zn limitation, analyzed by HPLC. Shown are averages of three independent replications.

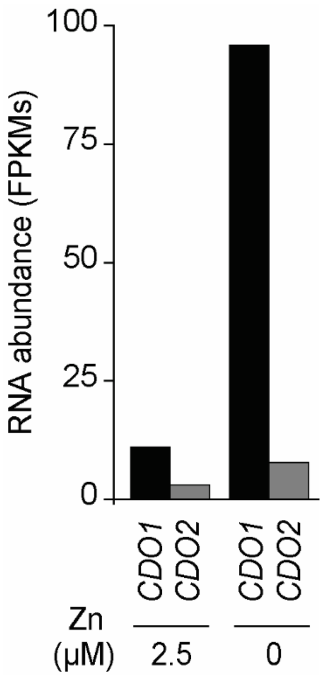


**Supplemental Figure 4. Increased expression of Cys oxygenases.** Shown are RNA abundance values determined by RNA-Seq in FPKMs of *CDO1 and CDO2* from cells grown either in Zn replete or Zn deplete medium as indicated.

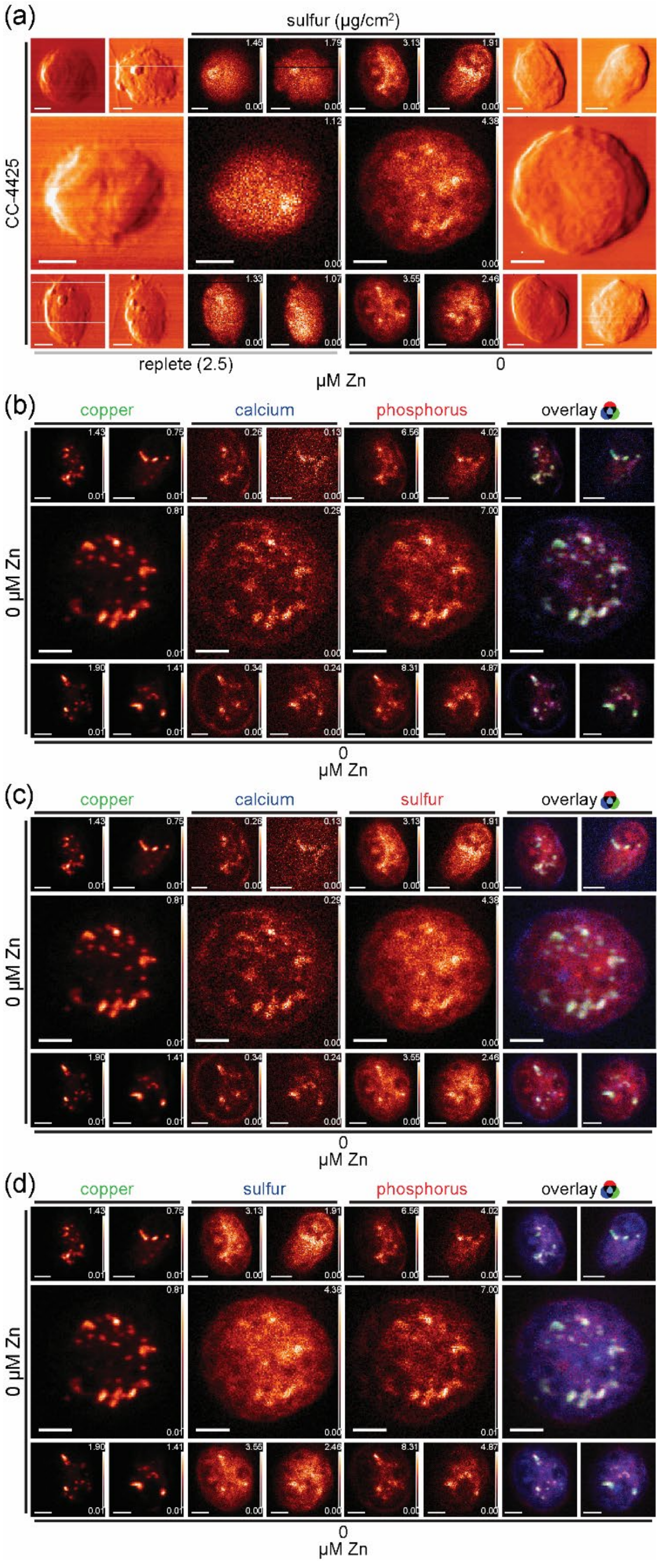


**Supplemental Figure 5. S is enriched at hot spots also enriched in C, P and Ca.**

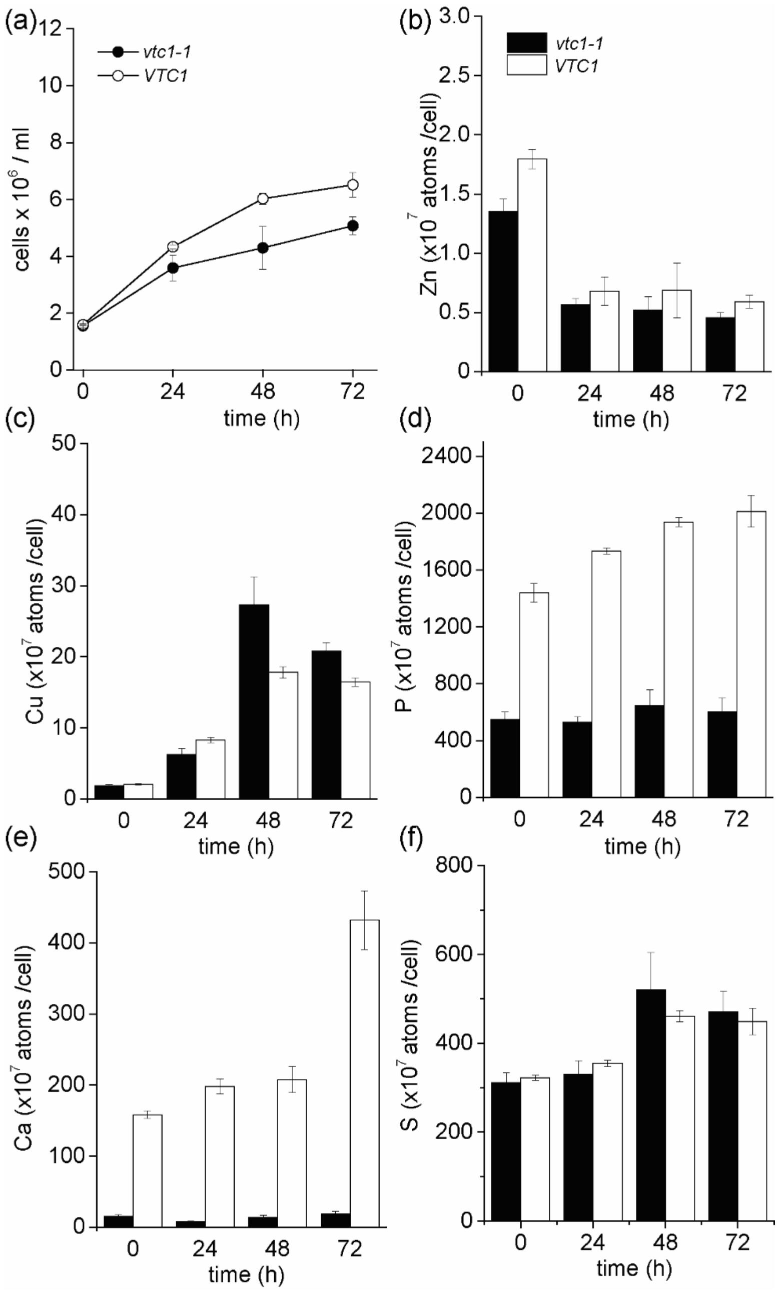


**Supplemental Figure 6. Cu accumulation is not affected in a Zn limited *vtc1* mutant.** *vtc1-1* mutants and complemented lines (*VTC1*) were grown in Zn limited medium, and 2x$10^6$ cells ml−1 were washed once with 1 mM EDTA and inoculated into medium lacking Zn. (a) Cells were counted on a coulter counter. Total Zn (b), Cu (c), P (d), Ca (e) and S (f) content was measured by ICP-MS/MS from three independent experimental replicates. Shown are averages and STDEV for each sample.

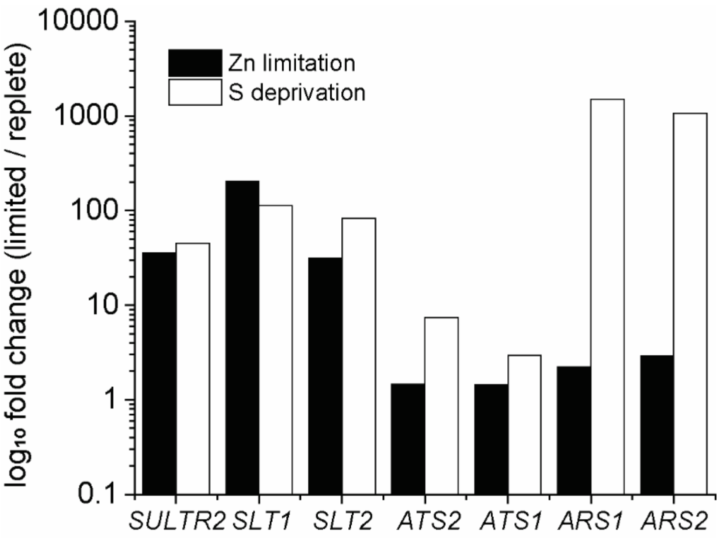


**Supplemental Figure 7. Overlapping and distinct gene expression patterns between S and Zn limitation.** Bar graphs depict differentially expressed genes in Zn limited and S deficient Chlamydomonas as indicated. Gene expression data are derived from experiments described in Hong-Hermesdorf et al. 2014 and González-Ballester et al. 2010.
